# Illumination of a progressive allosteric mechanism mediating the glycine receptor activation

**DOI:** 10.1101/2022.06.04.494798

**Authors:** Sophie Shi, Solène N Lefebvre, Laurie Peverini, Adrien Cerdan, Marc Gielen, Jean-Pierre Changeux, Marco Cecchini, Pierre-Jean Corringer

**Author notes:** equal contribution.

## Abstract

Pentameric ligand-gated ion channel mediate signal transduction at chemical synapses by transiting between resting and open states upon neurotransmitter binding. Here, we investigate the gating transition of the glycine receptor fluorescently labeled at the extracellular-transmembrane interface by voltage-clamp fluorimetry (VCF). Fluorescence reports a glycine-elicited conformational transition that precedes pore opening. Low concentrations of glycine, partial agonists or specific mixtures of glycine and strychnine trigger the full fluorescence signal while weakly activating the channel. Molecular dynamic simulations of a partial agonist bound-closed Cryo-EM structure show a highly dynamic personality: a marked structural flexibility at both the extracellular-transmembrane interface and the orthosteric site, generating docking properties that recapitulate VCF data. Data thus illuminate a progressive gating transition towards activation, displaying structural plasticity with novel implication concerning the mechanism of action of allosteric effectors.

## Introduction

Pentameric ligand-gated ion channels (pLGICs), including nAChRs, 5-HT_3_Rs, GlyRs and GABA_A_Rs constitute a superfamily of transmembrane receptors mediating intercellular communications in the nervous system ^1,2^. They transduce the binding of a neurotransmitter at their orthosteric site into the opening of an intrinsic ion channel, leading to ion fluxes that promote cell excitation or inhibition. The orthosteric binding site also binds partial agonists, with less efficacy than full agonists to elicit channel opening, and competitive antagonists. These distinct pharmacological profiles were interpreted in terms of the concerted two-state model (Monod et al., 1965) based upon a pre-existing equilibrium between the resting and active states that is differentially shifted depending on the nature of the effector ^3–5^.

In between these two equilibrium states, the conformational pathway that the protein takes during activation remain elusive. Single-channel recordings of numerous mutants of the muscle nAChR analyzed by REFERs (rate equilibrium linear free energy relationships) suggested an early motion of the extracellular domain (ECD) and a late motion of the transmembrane domain (TMD) during activation ^6^. Likewise, single-channel kinetics analysis on GlyRs and nAChRs were interpreted via a model involving a multistep process since accurate fitting of the experimental data require the introduction of intermediate states “flip”^7^ or “primed”^8^ in the gating transition. Although capturing structural reorganizations only indirectly, the electrophysiological data suggest that the transition is progressive, with the conformational changes starting from the ECD where the orthosteric site is located and propagating to the TMD to reach the channel gate. Moreover, computational studies of various pLGICs by Molecular Dynamics highlight their passage through a complex conformational landscape with the gating transition being composed of a progressive reorganization of the multimeric receptor architecture ^9–13^.

At the structural level, fluorescence experiments provided evidence that the bacterial pLGIC homologue GLIC undergoes a progressive reorganization, with a first pre-activation step involving a quaternary compaction of the ECD ^14^ followed by pore opening associated with a structural reorganization of the TMD and ECD-TMD interface ^15^. However, direct evidence for a progressive signal transduction in eukaryotic receptors is essentially lacking, despite recent cryo-EM structures of pLGICs revealing that the binding of orthosteric ligands is often accompanied by a significant reorganization of the extracellular domain (ECD), while the channel remains closed at the transmembrane domain (TMD), e.g. in 5-HT_3_R ^16,17^, GABA_A_R^18^, GlyR ^19^ and nAChR ^20^. Thus, pLGIC structures appear to be more flexible than previously anticipated, suggesting that ligands or classes of molecules could stabilize specific conformation(s) in the conformational landscape accessible to the protein. Whether the captured conformations solved by cryo-EM correspond or not to intermediates during transitions on the pathway to channel activation remains an open question. Addressing these questions requires monitoring the conformational dynamics of a fully functional receptor embedded in a plasma membrane and probing its relationship with both activation and desensitization. The Voltage-Clamp Fluorimetry (VCF) technique is perfectly suited to this purpose, since it allows the simultaneous recording of local conformational changes by fluorescence together with transmembrane current fluxes by electrophysiology.

Here, we used the ⍰1 homomeric GlyR, which is surely among the best characterized pLGIC by electrophysiology ^21,22^, cryo-EM alone or in complex with ligands ^19,16,23^, VCF ^24–31^, and MD simulations ^32,33^. We report the development and exploitation of a mutant GlyR⍰1 bearing a fluorescent sensor at the interface between ECD and TMD. Its pharmacological characterization using different allosteric effectors by VCF coupled to MD simulations and docking illuminates the occurrence of intermediates, displaying dynamic properties of the ECD, on the path to channel opening consistent with a progressive mechanism of gating.

## Results

### 1 Design of a fluorescent-quenching sensor reporting conformational changes at the ECD-TMD interface

To generate a fluorescent sensor related to the activation process, we compared the structure of the zebrafish GlyR⍰1 in the resting-like apo state (PDB:6PXD) with the glycine-bound open-channel structure (PDB:6PM6) ^19^. Beside the well documented contraction of the orthosteric site at the subunit-subunit interfaces in the ECD and the opening of the channel in the TMD, visual inspection also shows an expansion of the ECD-TMD interface, where Cys-loop of each subunit moves away by nearly 3 Å from the pre-M1 region of the adjacent subunit (Figure 1A, B). In order to monitor this reorganization, we used the tryptophan quenching technique that can sense the relative place of two residues ^34,35^. This technique is particularly relevant to monitor distance changes of 5-15 Å range ^14,15^. We introduced a cysteine at position Q219 in the pre-M1 region previously used for labeling with fluorescent dyes ^28^, together with a tryptophan in Cys-loop at position K143, in order to quench the fluorophore in a conformation dependent manner (Figure 1 A,B).

**Figure 1.**
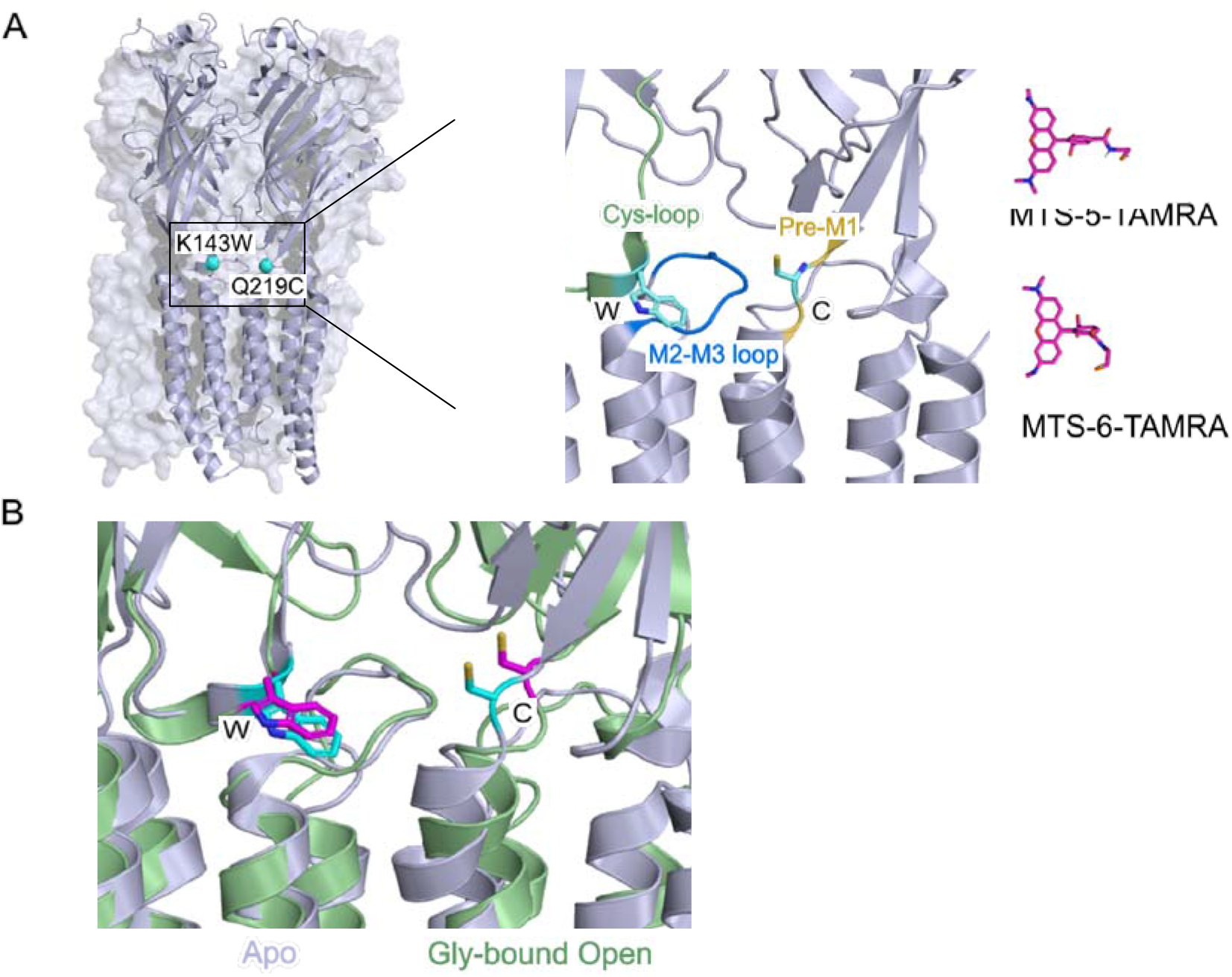
Localization of K143W/Q219C sensor in the ⍰1 glycine receptor structure. **(A)** Side view of the zebrafish ⍰1 glycine receptor structure in Apo state (PDB:6PXD) showing the position of the mutated residues K143W in Cys-loop and Q219C in Pre-M1 loop. Right panel: representation of the two isomers of 5(6)-carboxytetramethylrhodamine methanethiosulfonate (MTS-TAMRA) used for the labeling of the mutated cysteine. **(B)** Structural comparison of the quenching sensor K143W/Q219C between the Apo state in grey (PDB:6PXD) and Gly-bound open state in green (PBD:6PM6), residues are presented in cyan for apo state and magenta for Gly-bound open state. The distance calculated between the C⍰ of the two residues varied from 12,9 Å for Apo state to 15,6 Å for Open state.

### 2 The quenching sensor K143W/Q219C reveals glycine-elicited fluorescence variations at much lower glycine concentrations than currents

The endogenous extracellular single cysteine of the human GlyR⍰1 was mutated to avoid non-specific labeling, yielding WT-C41V displaying wild-type like properties in electrophysiology (Figure S1). Upon introduction of the quenching sensor mutations, the unlabeled K143W/Q219C/C41V was analyzed by two-electrode voltage clamp electrophysiology (TEVC). In all dose-response curves presented here, we used a brief perfusion (< 10 seconds) of the glycine solutions to reach a current plateau with no apparent desensitization. The unlabeled K143W/Q219C/C41V displays an EC_50_^current^ that is 34-fold higher than that of WT-C41V (Figure 2 A, B & S1, Table 1), indicating a loss-of-function phenotype. We controlled that the mutations do not impair ion channel permeation by performing outside-out single-channel recordings (Figure 2C) and measuring a unitary conductance of 86.7 ± 4.6 pS that is identical to that of WT-C41V (85.7 ± 4.5 pS) (Figure S1). After labeling with MTS-TAMRA (Figure 1A), K143W/Q219C/C41V was analyzed by VCF, using a dedicated recording chamber (Figure S2). Its EC_50_^current^decreases by 1.6-fold, indicating that the labeling produces a slight gain-of-function that partially counterbalances the effect of the K143W/Q219C mutations (Figure S3A). Robust glycine-elicited fluorescence variations were also observed (Figure 2A, D). They start at non-activating glycine concentrations (5-25 μM) and reach a plateau at the beginning of the current dose-response curve, with a maximal fluorescence amplitude ΔF/F corresponding to a 12% decrease (Figure 2A). Therefore, the EC_50_^fluo^ is 73-fold lower than the EC_50_^current^ (Figure 2B). As a control, we also recorded the labeled Q219C/C41V. It shows only maximal fluorescence amplitude ΔF/F corresponding to a 2% decrease (Figure S3B), indicating that the fluorescence variation of the K143W/Q219C sensor is dominated by the quenching provided by the introduced tryptophan. Of note, the MTS-TAMRA used here corresponds to a commercial mixture of isomers (Figure 1A), each of which was verified to produce the same VCF pattern separately (Figure S3C).

**Figure 2.**
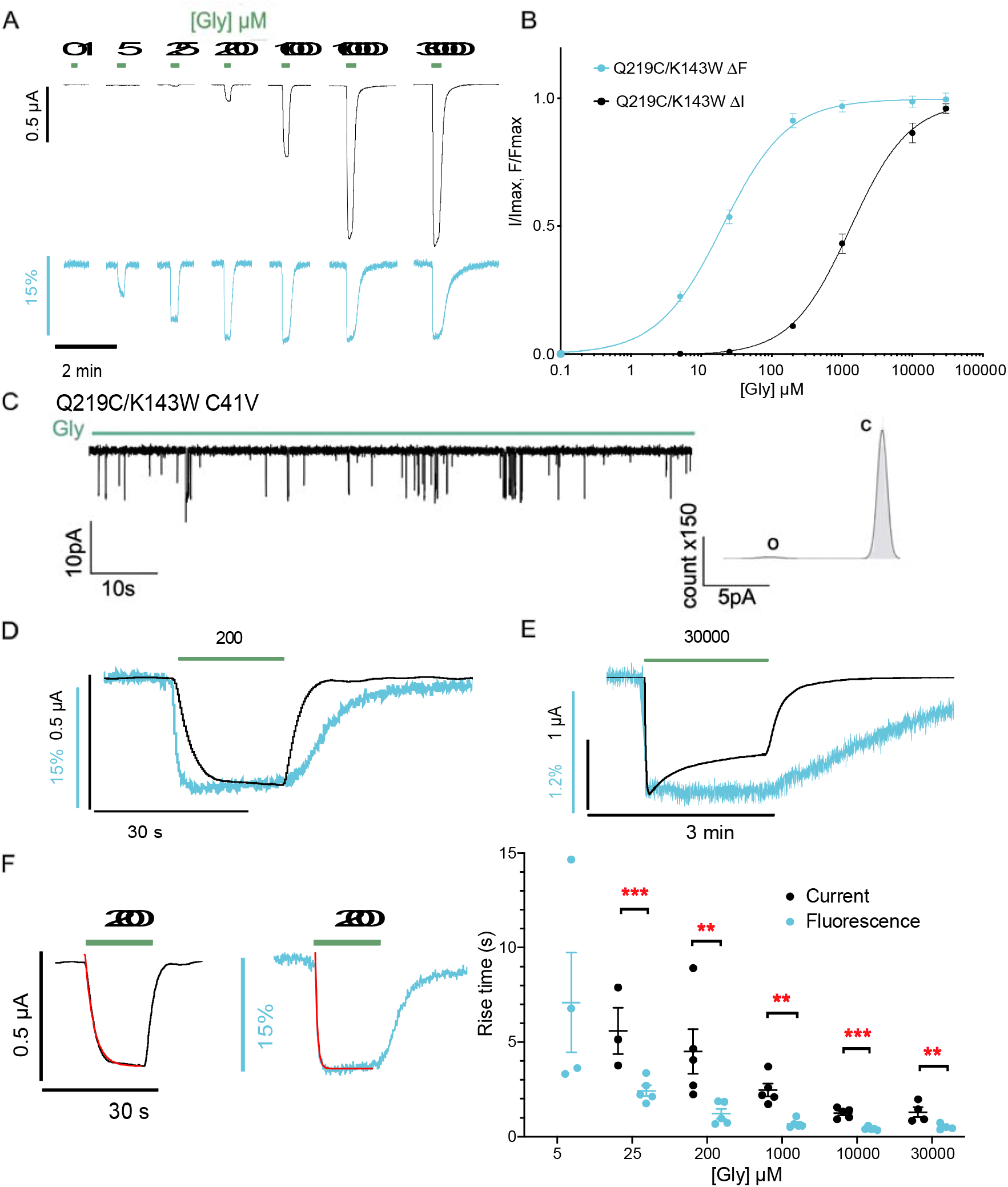
Electrophysiological and fluorescence characterization of K143W/Q219C/C41V by VCF. **(A)** Representative VCF recordings on oocytes of the mutant labeled with MTS-TAMRA (black for current and cyan for fluorescence). Glycine application triggers a current variation and a fluorescence quenching phenomenon with a maximum of fluorescence variation that reach 12.1 ± 1.1 % of ΔF/F. At low concentration of glycine (under 25 μM), only fluorescence quenching is observed without any current. **(B)**ΔI (black) and ΔF (cyan) dose-response curves with mean ± S.E.M. values (n=5). **(C)** Left panel: representative trace of single-channel recordings obtained in outside-out configuration recorded at −100mV with concentrations of glycine at 100μM. Right panel: histograms of current amplitude representing the closed state (c) and the open state (o). **(D)** Superimposition of the current and fluorescence recordings evoked by 200 μM glycine application showing that the onset of the fluorescence is faster than the onset of current and that the fluorescence offset is slower than the current offset. **(E)** Representative recording with a high glycine concentration (30mM) application showing that the desensitization triggered by a prolonged glycine application does not impact the fluorescence variation (n=4). **(F)** Right panel: single exponential fitting (red line) of the current (right) and fluorescence (left) traces onset. Left panel: time constants τ (onset) values obtained via single exponential fitting with mean and S.E.M. error bars in second at different glycine concentrations.); Unpaired student t-test revealed the significance of the difference between fluorescence and current onset (**: *P* < 0,005; ***: *P* < 0,0005).

**Table 1.** EC_50_ values for current and fluorescence responses to glycine at labeled and unlabeled GlyR⍹1 WT and mutants.

|  | EC<sub>50</sub><sup>current</sup> (μM) | n<sub>H</sub> | EC<sub>50</sub><sup>fluo</sup> (μM) | n<sub>H</sub> | n |
| --- | --- | --- | --- | --- | --- |
| WT | 165.84 ± 14.06 | 1.55 ± 0.08 |  |  | 5 |
| WT unlabeled | 79.30 ± 11.34 | 2.16 ± 0.36 |  |  | 8 |
| WT C41V | 162.87 ± 43.21 | 1.18 ± 0.13 |  |  | 5 |
| WT C41V unlabeled | 152.40 ± 28.87 | 1.33 ± 0.26 |  |  | 4 |
| Q219C C41V | 56.80 ± 6.65 | 1.84 ± 0.13 | 550.95 ± 67.55 | 1.71 ± 0.15 | 6 |
| Q219C C41V unlabeled | 70.52 ± 3.47 | 2.23 ± 0.46 |  |  | 5 |
| Q219C/K143W C41V | 1541.12 ± 338.71 | 1.06 ± 0.11 | 21.00 ± 2.26 | 0.96 ± 0.10 | 5 |
| Q219C/K143W C41V unlabeled | 2485.20 ± 145.08 | 1.04 ± 0.03 |  |  | 5 |
| Q219C/K143W/N46D/N61D C41V | 23029.40 ± 4281.72 | 1.41 ± 0.04 | 1428.52 ± 396.81 | 1.12 ± 0.06 | 5 |

### 3 The quenching sensor K143W/Q219C reports an “intermediate” conformational reorganization occurring before the onset of currents

Inspection of the current/fluorescence time courses and quantification by single exponential fitting reveals important features. First, the onset of both the fluorescence and current traces is faster with increased glycine concentrations whereas their offset during agonist rinsing is slower (Figure 2F, Table S3, Figure S4). Second, at each glycine concentration, the rise time of the fluorescence change is systematically faster than that of the currents (Figure 2F) whereas the offset tends to be slower particularly at glycine concentrations >200 μM (Figure S4). Noteworthy, these kinetic values are limited by the rate of agonist perfusion in the recording chamber, and do not reflect the intrinsic kinetics of the receptor (Figure S3). For instance, upon perfusion of 200 μM Gly (Figure 2D), the glycine concentration at the oocyte surface will progressively rise, first reaching the concentration eliciting full fluorescence variation (100 μM), then reaching after a delay the concentration eliciting currents (200 μM). Conversely, upon glycine washing, its concentration will rapidly drop below 100 μM, causing deactivation, but will take more time to drop below the concentrations causing fluorescence variations. The relatively slow kinetics of perfusion of the agonist thus separate in time the molecular events causing the fluorescence variation and the current, the former preceding the later.

Figure 2 A, B shows that 100 μM glycine, at steady state, elicits most of the fluorescence variation but no significant current. This indicates that the receptor has completed the full movement reported by the quenching sensor with a closed channel. Hence, VCF data provide direct evidence for a conformational transition toward one state or a cluster of states that are structurally distinct from both resting and active. Importantly, kinetic data show that these conformations appear before channel activation in our recording conditions (Figure 2D, F). This suggests that they correspond to early “intermediate conformation(s)” along the allosteric transition pathway to activation that are not related to desensitization. To further investigate this possibility, we applied glycine at high concentration (30 mM) for a prolonged period (1 min), leading to a rapid onset of the current followed by a slower decrease (34.6 ± 4.9 % decrease after 1 min) caused by desensitization. The fluorescence strongly decreases during the activation phase, then reaches a plateau and remains stable during the second phase, providing evidence that the quenching sensor does not report movements related to desensitization (Figure 2E).

### 4 The Gly-elicited fluorescent and current changes are linked to glycine binding to the orthosteric site

The marked separation of the current (ΔI) and fluorescence (ΔF) dose-response curves raises the possibility that both processes could be mediated by different classes of glycine binding sites within and outside the orthosteric site. To challenge this possibility, we introduced two mutations into the orthosteric binding site (N61D/N46D, Figure 3A) that strongly decrease the affinity for glycine ^36^. The K143W/Q219C/N61D/N46D/C41V shows a parallel rightward shift of both ΔI and ΔF curves (15-fold and 68-fold increase in EC_50_^current^ and EC_50_^fluo^, respectively) (Figure 3B, C, Table 1), indicating that both processes are strongly linked to glycine binding to the orthosteric site.

**Figure 3.**
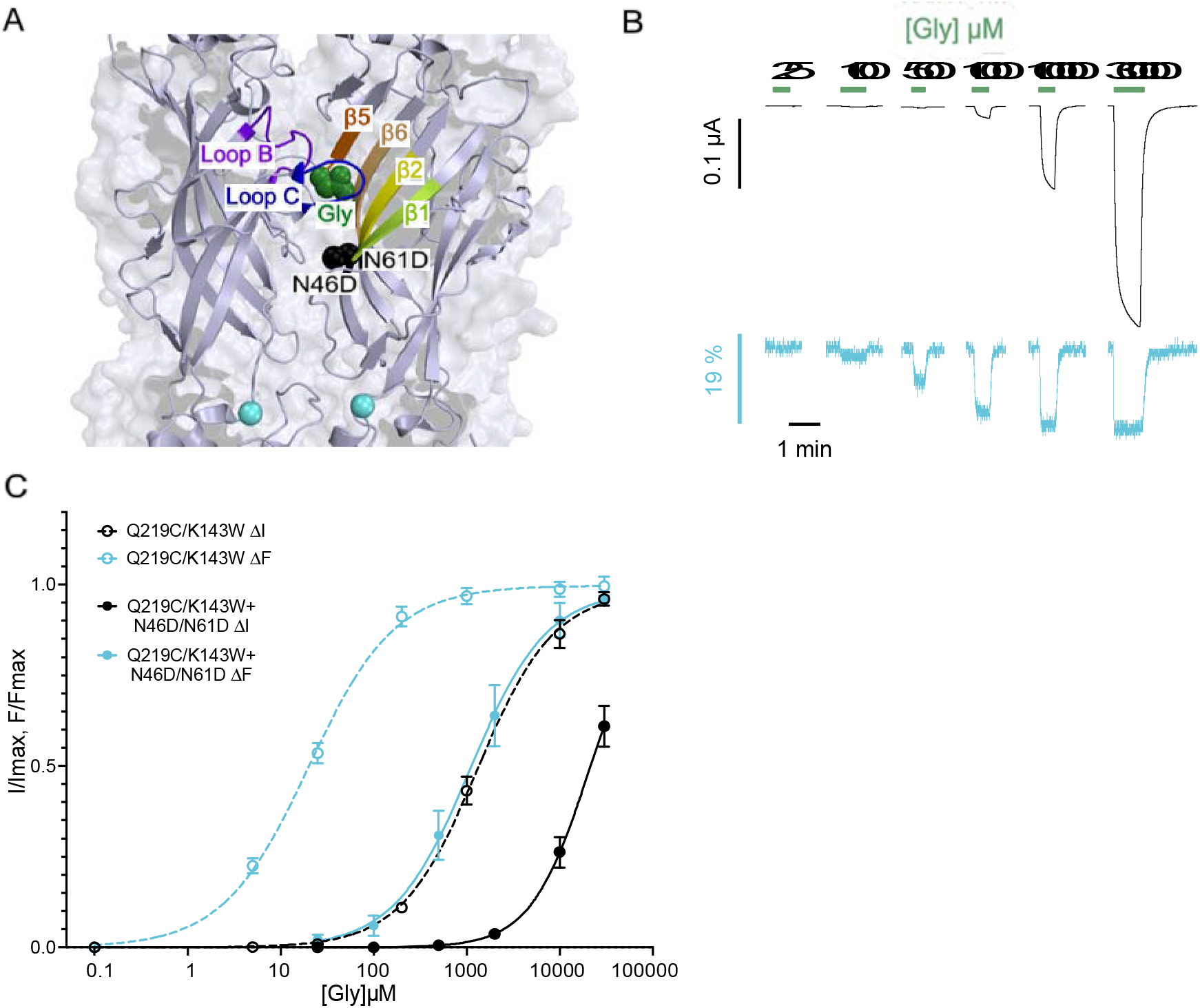
Effect of N46D/N61D/C41V mutations on ΔI and ΔF curves of K143W/Q219C. **(A)** Side view of zebrafish ⍰1 glycine receptor structure in Apo state (PDB:6PXD) with details of loops and β-sheets forming the orthosteric site and the localization of N46D/N61D mutations. Glycine molecule is represented in green and N46D/N61D mutated residues (spheres) in black. **(B)** Representative VCF recordings of the mutant K143W/Q219C + N46D/N61D (current in black and fluorescence in cyan). **(C)** ΔI (cyan) and ΔF (black) dose-response curves with mean ± S.E.M. show a parallel shift of the current and fluorescence curves of the K143W/Q219C/N46D/N64D/C41V mutant (full line; n=5) compared to K143W/Q219C/C41V alone (dotted line).

### 5 Partial agonists, strychnine and propofol differentially affect the current and fluorescence variations

First, we investigated the effect of agonists that bind at the orthosteric site but are less effective than glycine at activating the receptor, i.e., partial agonists. We selected β-alanine and taurine: the first one triggers 54-98% of the glycine-elicited maximum current ^37–39^ and the second one leads to 20-42% of the glycine-elicited maximum current, all measured in oocytes by TEVC ^37,40,41,39^. On K143W/Q219C/C41V, we found that their efficacy decreases, eliciting only 6.1 ± 2.2 % and 2.0 ± 1.4% of the glycine-elicited maximum current respectively (Figure 4). Such a decrease in efficacy is commonly observed for loss-of-function mutants ^31^.

**Figure 4.**
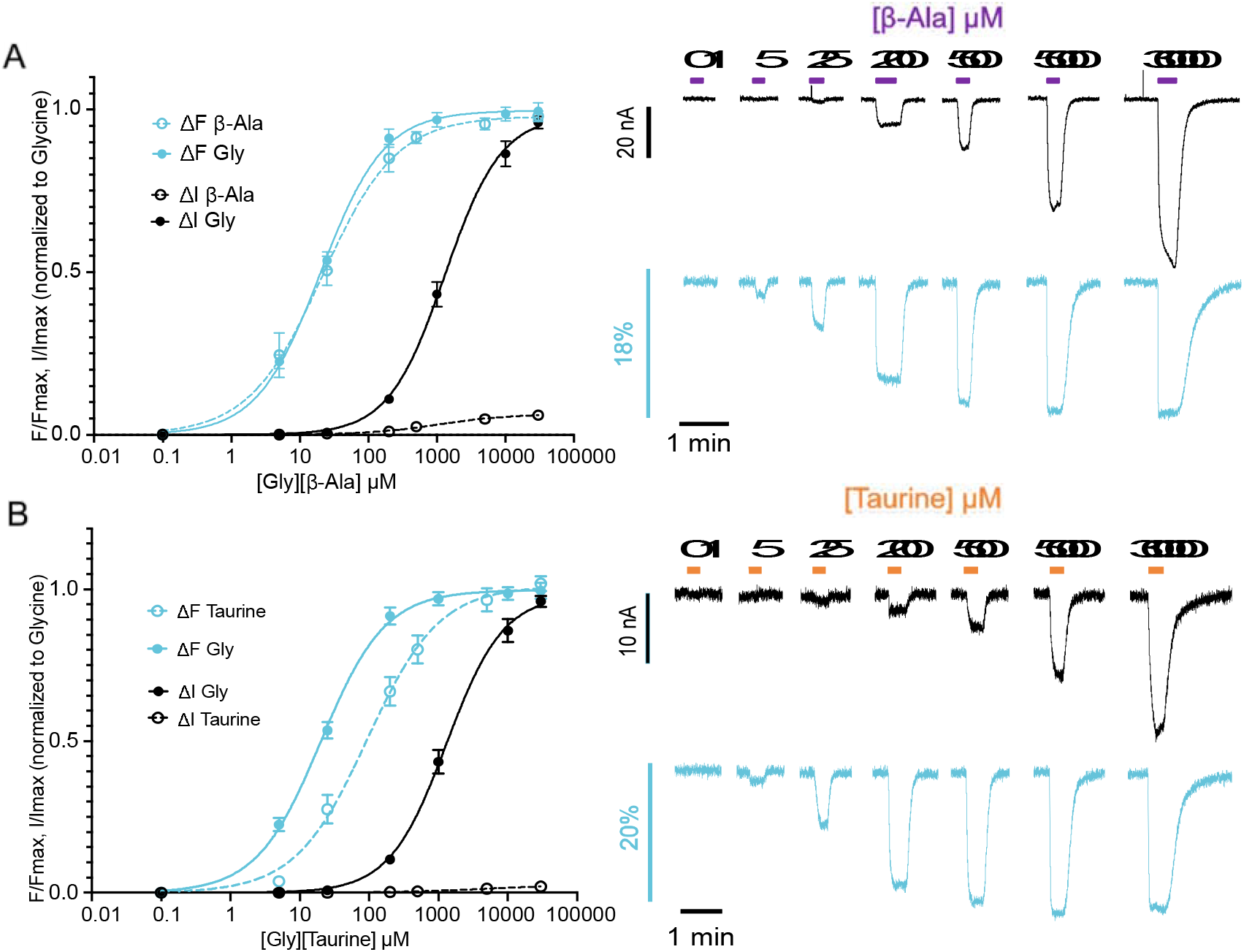
Electrophysiological and fluorescence characterization of partial agonist (β-alanine and taurine) effects on K143W/Q219C/C41V. **(A)** Left panels: the ΔI (cyan) and ΔF (black) dose response curves (normalized to the glycine maximal response recorded on each individual oocyte) with mean and S.E.M. for K143W/Q219C under glycine (full line) and β-alanine (dotted line) application. The fluorescence dose-responses curves for both molecules are superimposed illustrating that the efficacy of the β-alanine to activate the receptor is lower than the glycine (only 6.1 ± 2.2 % of glycine maximum response in current, n=6). Right panel: representative VCF recording on oocytes under β-alanine application. **(B)** Left panel: ΔI (cyan) and ΔF (black) dose response curves (normalized to the glycine maximal response recorded on each individual oocyte) with mean and S.E.M. for K143W/Q219C under glycine (full line) and taurine (dotted line) application. Taurine elicits only 2.0 ± 1.4 % of glycine maximum current, n=6. Right panel: representative VCF recording in oocytes under taurine application.

Surprisingly, both partial agonists elicit the full fluorescence variation recorded with glycine (Figure 4A, B). β-Alanine and taurine also show a large difference in the fluorescence and current dose-response curve, their EC_50_^fluo^ being respectively 66- and 70-fold lower than their EC_50_^current^ (Table 1). β-Alanine and taurine thus trigger the full transition mediating the fluorescence change, but they are very weakly effective to trigger the “downward” process of activation.

Next, we investigated the competitive antagonist strychnine. When strychnine is applied alone at 5 μM, no current is observed as expected, but the fluorescence increases over the baseline in an opposite direction to the glycine-elicited quenching (dequenching) and reaches a plateau corresponding to 30-40% of the maximal glycine elicited fluorescence response. We then applied strychnine (5 μM) during a perfusion of glycine at different concentrations (Figure 5). At 100 μM glycine, strychnine inhibits both the current and fluorescence variations elicited by glycine. As before, strychnine tends to increase the fluorescence over the baseline, but the effect is not significant. At 1 mM of glycine, strychnine still totally inhibits the current, but surprisingly the glycine-elicited fluorescence quenching is decreased by only 48%. Finally, at 10 mM of glycine, strychnine inhibits only 77% of the current, while the fluorescence is almost not affected. Therefore, co-application of strychnine with various concentrations of glycine differentially affects the fluorescence variation and gating currents.

**Figure 5.**
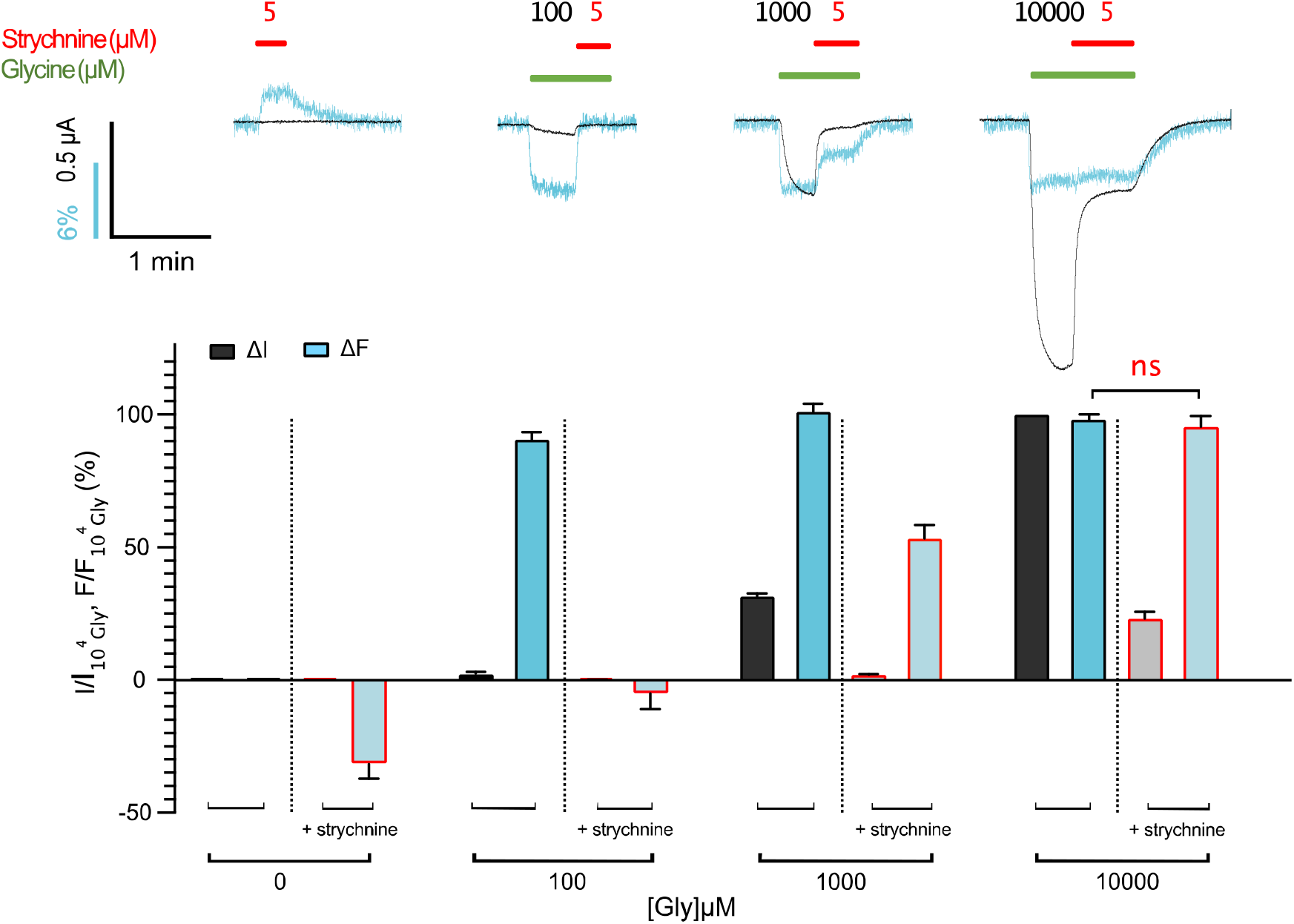
Differential inhibition of strychnine on fluorescence and current on K143W/Q219C/C41V. Upper panel: representative VCF recording superimposing the current (black) and fluorescence (cyan). left panel: application of strychnine alone at 5μM triggers a fluorescence dequenching, right panels: glycine is applied at different concentrations first alone (green bar) and then in mixture with 5μM strychnine (red bar plus green bar), showing differential strychnine-elicited inhibition of current and fluorescence depending on the glycine concentration. Lower panel: fluorescence (cyan) and current (black) variations normalized to the fluorescence and current variations under 10mM of glycine (mean and S.E.M. (n=5)). “ns” Denotes not significantly different between the ΔF (under glycine) and ΔF (under strychnine inhibition); *P* < 0.05 (Unpaired student t-test).

Last, we investigated the general anesthetic propofol, a positive allosteric modulator of GlyR⍰1 that binds in the TMD of pLGICs ^42–45^. At concentrations below 300 μM, propofol elicits a fluorescence variation of about 3-5% ΔF/F without activating the receptor, while at higher concentrations it also activates the currents in the absence of glycine (Figure S5).

In conclusion, pharmacological analysis shows that the intermediate conformation(s) is stabilized in conditions yielding no significant activation: saturating concentrations of partial agonists, a specific range of strychnine plus glycine mixtures, and non-activating concentrations of propofol.

### 6 Molecular dynamics simulations of a cryo-EM partial-agonist bound state recapitulates the pharmacological profile of the intermediate state(s)

A very recent cryo-EM analysis of the zebrafish GlyR ^19^ has revealed that the partial agonists taurine and GABA may stabilize a previously unseen closed-channel state (referred to as “tau-closed”) characterized by a significant reorganization of the ECD compared to the apo-state. We hypothesized that the intermediate conformation(s) illuminated by VCF is structurally similar to tau-closed. To explore this hypothesis, we carried out all-atom MD simulations of the apo-closed (PDB:6PXD), tau-closed (PDB:6PM3) and tau-open (PDB:6PM2) structures embedded in a POPC bilayer. In addition, since strychnine nicely fits the orthosteric site of apo-closed, we performed MD simulations of the strychnine-closed (stry-closed) complex as well. For each system, five independent simulations of 100 ns were carried out.

The structural stability of the protein was analyzed by monitoring the root-mean-square fluctuations (RMSF) from the average structure extracted from MD (Figure 6A). In addition, we calculated the evolution of the twisting and blooming angles of the ECD during each simulation (Figure S6). The simulations show that: i. apo-closed is the most flexible conformational state of the receptor (Figure 6A) and strychnine binding stabilizes the bloomed conformation (separation of the ECD seen in the Apo-closed cryo-EM structure), rigidifying the ECD; ii. the ECD of tau-closed is remarkably more flexible than that in tau-open despite the apparent structural similarity in the cryo-EM coordinates that show an un-bloomed (compaction of the ECDs) structure; iii. tau-closed has a weaker affinity for taurine than tau-open as evidenced by four spontaneous unbinding events in the tau-closed simulations, that are associated with significant blooming of the structure; iv. the C-loop that is critical for orthosteric ligand binding is more flexible in tau-closed (RMSF of 3.5 Å) than in tau-open (RMSF of 2.5 Å) or stry-closed (RMSF of 2.0Å); v. the conformational dynamics of the ECD-TMD interface (i.e., β1-β2 loop, Cys-loop, and the β8-β9 loop) is enhanced in tau-closed relative to apo-closed and tau-open. Taken together, the simulation results highlight that the tau-closed conformation features a surprising dynamic character, which is remarkably different from that of tau-open and apo-closed and could not be deduced from the comparison of static cryo-EM structures only (Figure 6A).

**Figure 6.**
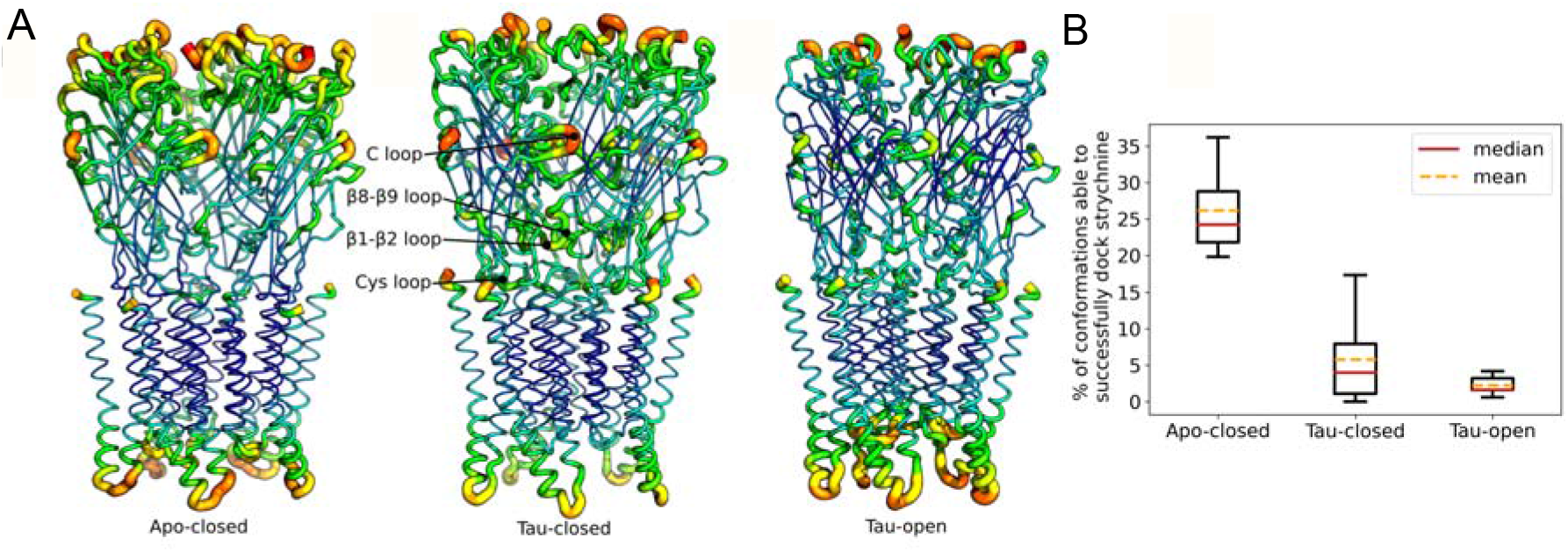
Dynamic “personality” of the tau-closed state. (**A**). Root-mean-square fluctuations (RMSF) from the average structure extracted from 0.5 μs MD simulations in explicit solvent/membrane are shown for three conformational states of GlyR captured by cryo-EM. Apo-closed corresponds to the resting state, tau-open to the active state of the receptor. For each state, RMSF values ranging from 0.5 Å to 4.5 Å are shown by the color (from blue to red) and the thickness (from thin to thick) of the sausage representation (**B**). State-based docking of strychnine. Relative strychnine-binding affinities for apo-closed, tau-closed, and tau-open was probed by docking strychnine to an ensemble of 500 protein snapshots sampled by MD and comparing the success rate of docking; i.e. a docking experiment was considered as successful if the docking score was within 10% of the score obtained in the strychnine-bound X-ray structure (stry-closed). The mean (yellow) or median (red) dashed lines show that strychnine binding in tau-closed is significantly stronger than in tau-open.

To explore the correlation between MD and VCF data, we quantified the reorganization of the fluorescence quenching sensor, as well as the binding of strychnine.

First, we calculated the average distance between the Cα carbons of the VCF quenching sensor (corresponding to K143W/Q219C). It is 13.0 ± 0.7 Å in stry-closed and apo-closed and this value increases to 14.7 ± 1.4 Å and 16.4 ± 1.0 Å in tau-closed and tau-open, respectively. Therefore, the significant reorganization seen in the simulations of the sensor from the apo to the tau-closed conformation is compatible with a change in fluorescence during this transition. We speculate that, during the separation of the Cα carbons, the rather long and flexible TAMRA “side chain” linked to C219 reorganizes to interact more frequently with W143, generating fluorescence quenching.

Second, the binding of strychnine to the various conformations of GlyR was explored by docking. As expected, favorable docking scores were obtained in stry-closed (−12.1 kcal/mol) and apo-closed (−10.3 kcal/mol) from cryo-EM. By contrast, both tau-open and tau-closed from cryo-EM feature an orthosteric site that is too small to accommodate strychnine and the docking scores were largely unfavorable. To account for the intrinsic flexibility of the protein, the docking experiments were repeated using protein conformations extracted from the MD trajectories. Assuming that a docking experiment was successful when the docking score was < −9.3 kcal/mol (i.e. within 10% of the score obtained in stry-closed), the success rate for docking strychnine was determined by counting the fraction of successful docking over 500 protein snapshots from MD of apo-closed, tau-closed, and tau-open. As shown in Figure 6B, docking of strychnine was successful at 26% (std dev 6.5%) in apo-closed, 6.6% (std dev 7.0%) and with large deviations among replicas in tau-closed, and only 2.2% (std dev 1.4%) in tau-open. We conclude that the enhanced flexibility of the tau-closed structure makes it very different from tau-open and compatible with strychnine binding.

Hence, the simulation results match with the VCF data presented above and support the conclusion that the intermediate conformation(s) revealed by fluorescence is consistent with a highly dynamic cluster of states simulated from the tau-closed structure captured by cryo-EM.

## Discussion

### 1 Structural evidence for an early intermediate step in the activation path revealed by VCF

The VCF data upon glycine application reveals an intermediate conformation(s) characterized by a change in fluorescence but no current, followed by channel opening with no further change in fluorescence. Three lines of evidence support that the intermediate conformation(s) illuminated by VCF is on-path to activation: i. Receptors in this intermediate conformation(s) are clearly activatable since they appear before the onset of the currents, and are thus not desensitized, ii. Receptors in the intermediate conformation(s) are stabilized by partial agonists, low glycine concentrations, and mixtures of glycine and strychnine, showing a pharmacological profile “in between” those stabilizing the resting and active states, and iii. MD simulations of several cryo-EM structures support the conclusion that the intermediate conformation(s) by VCF is consistent with a structural reorganization of the resting state where activation of the ECD is underway, while that of the TMD is not. Of note, the mutations performed here to introduce the fluorescence quenching sensor cause a significant loss-of-function phenotype characterized by a 10-fold increase in EC_50_^current^ of glycine, and a decreased efficacy of the partial agonists to activate the receptor, yet with an intact conductance of the channel. It is thus likely that, in the WT context, a lower ratio of the intermediate(s) to active state would occur in the presence of partial agonist, as suggested by cryo-EM data in the presence of taurine ^19^. In addition, we found that application of strychnine alone produces an opposite fluorescence variation compared to agonists. This suggests that strychnine alone stabilizes the receptor in a conformation distinct from the apo state, or alternatively that the intermediate conformation(s) are already significantly populated in the absence of ligands and strychnine shifts the equilibrium toward the resting state ^46^.

Previous single-channel works on nAChRs and GlyRs suggested the contribution of flipped/primed intermediate states on receptor’s activation. Flipped/primed are incompletely stabilized by partial agonists, underlying their limited efficacy to fully activate the receptor ^8,47^. This is in contrast with the intermediate conformation(s) unraveled here, which is fully stabilized by partial agonists and consequently must occur earlier than the flip/prime transition.

### 2 MD simulations of the Cryo-EM “Tau-closed” state provides a molecular basis for the VCF data

An important observation from VCF is that taurine binding stabilizes primarily the intermediate state(s). The structure of the “tau-closed” state of the zebrafish GlyR recently solved by cryo-EM is a strong candidate for an early on-pathway intermediate during activation ^19^. However, docking experiments show that this structure is unable to accommodate the bulky antagonist strychnine, a feature that is inconsistent with our VCF data. Interestingly, MD simulations of the tau-closed structure provide a way to explain this discrepancy, revealing that its ECD is surprisingly dynamic in the tau-closed structure and samples the spontaneous opening of the orthosteric site on the sub-μs time scale, which makes it compatible with strychnine binding. Combined with the VCF data, these observations suggest that receptors in presence of strychnine and agonists mixtures would keep an overall intermediate-like conformation with significant asymmetry at the orthosteric sites, some bound to agonists/partial agonists in an active-like compact state, while others bound to strychnine in a resting-like expanded state. We speculate that this mechanism also applies in the case of partial occupation of the orthosteric sites by glycine, accounting for the VCF data at low glycine concentration and of the large separation of the ΔI and ΔF dose-response curves.

It is noteworthy that the simulations reveal that the tau-closed state presents a unique dynamic personality that is remarkably different from those at the endpoints of activation. In fact, comparison of the atomic fluctuations in the different conformations of GlyR reveals that the flexibility of the ECD-TMD interface is surprisingly enhanced in tau-closed relative to both apo-closed (resting-like) and tau-open (active-like). We speculate that the unique dynamic signature of tau-closed originates from the hybrid nature of this state, which features an ECD in a nearly active conformation along with a resting-like TMD endowed with a closed channel. Furthermore, the apparent mismatch at the ECD-TMD interface, which enhances the flexibility of the ECD domain and its blooming in tau-closed, favorizes the structural plasticity for ligand binding. It leads the receptor to undergo intermediate state(s) in between resting and active, where the conformational transition of the ECD is essentially complete, while that of the TMD has not yet occurred. This hypothesis is consistent with the large re-organization of the pre-M1 and Cys-loop regions probed by VCF in the absence of currents and suggests that the tau-closed structure and dynamics provide a reasonable representation of the early intermediate conformation.

From combined VCF and MD data, we thus propose a speculative model of receptor activation depicted in Figure 7, where the receptor transits through a cluster of highly dynamic intermediate states, allowing significant local asymmetry at the level of the orthosteric site, but a more symmetric transition at the ECD-TMD interface where all fluorescent sensors reorganize.

**Figure 7.**
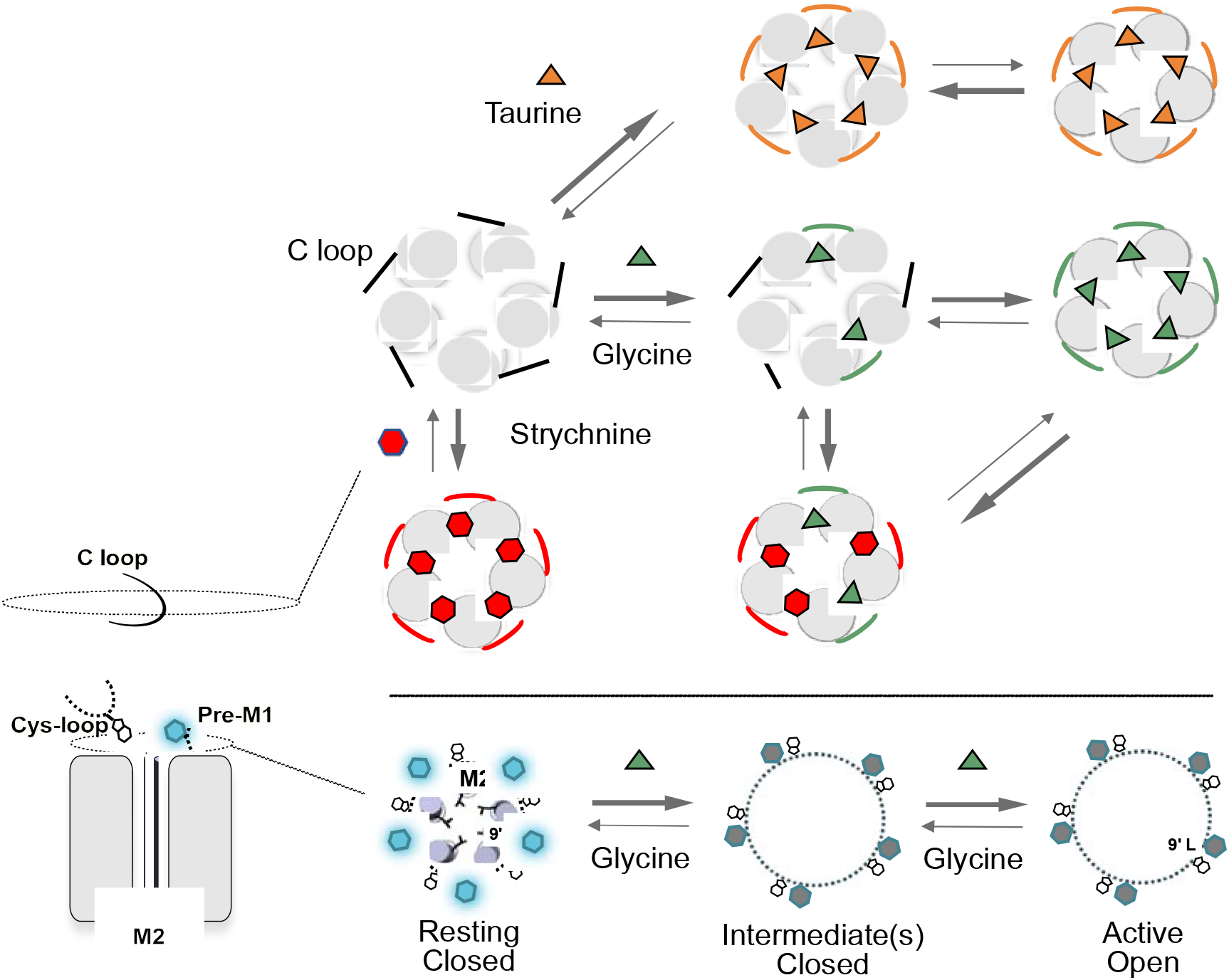
Hypothetical transition pathway of the GlyR. Left panel: side-view cartoon representation of two subunits of the GlyR. C loop contributing to the orthosteric site, the quenching sensor, the TAMRA (hexagon) and tryptophane (indole) are represented. Upper panel: top view representation of the receptor in different states with the ECD of each subunit shown in circle and C loop as a line. Lower panel: top view representation of the TMD showing the M2 helixes and the L9’ that closes the pore in the resting and intermediate states. The black circle represents positions where the fluorescent quenching sensor is grafted. The indole goes closer to the TAMRA from the resting to the intermediate(s) states, generating a quenching of fluorescence.

### 3 About the occurrence of a progressive multistep transition in other pLGICs

Among pLGICs, the GlyR has been extensively studied by VCF by Lynch *et al,* labeling the upper M2 helix at position 19’^25,31^, the orthosteric site, loop 2 and pre-M1 regions ^28,30^, and the TMD ^26,27^. In all cases, the fluorescence dose-response curves were superimposed or right-shifted as compared to the current dose-response curve. Thus, the phenotype of the quenching sensor observed here, which is characterized by a major leftward shift of the fluorescence dose-response curve, is unique and reports previously unseen molecular motions. Interestingly, phenotypes resembling the one presented here have been already described for the bacterial channel GLIC and other cationic pLGICs. With GLIC, we reported a “pre-activation” phenotype for a series of quenching sensors at the ECD interface and M2-M3 loop ^14,15^. They all report fluorescence variations at non-activating agonist (proton) concentrations and the fluorescence variation was faster (ms range measured by stopped-flow) than the onset of activation (10-100 of ms measured by patch-clamp) ^48^. It is noteworthy that M2-M3 loop is located just in between the quencher (W143) and the fluorophore (C219) in the GlyR structure (Figure 1).

In case of cationic pLGICs using VCF, labeling of the muscle nAChR (αγαδβ subtype) at the top of TMD (position 19’ located at the upper end of M2 helix and facing the ECD) reports a 100-fold lower EC_50_^fluo^ as compared to EC_50_^current^ for Ach (binding the αδ interface), while no difference was observed for epibatidine (binding the αγ interface). Moreover, the change in fluorescence occurs with fast kinetics and pharmacological considerations support the occurrence of a conformational transition toward a singly liganded closed-channel state ^49^. Labeling at the 19’ position has also been performed on the 5-HT_3_R, consistent with monitoring a pre-active reorganization ^17^. Finally, labeling of the 04β4 receptor in the extracellular loop 5 of ⍰4 shows a 5-fold lower EC_50_^fluo^ as compared to EC_50_^current^ for ACh suggesting that loop 5 moves before channel activation. Additionally, the antagonist DHβE elicits also a robust fluorescent change ^50^ which suggests a highly dynamic nature of loop 5 in the activation and inhibition process.

Altogether, data on the GlyR, GLIC and possibly nAChR/5HT_3_R provide evidence for intermediate transitions involving structural reorganizations mainly at the ECD and/or ECD-TMD interface. This idea is in line with REFERs analysis of the muscle nAChR, that suggests that the orthosteric site and M2-M3 loop move very early during the activation transition ^6^. It is also consistent with simulations describing the gating isomerization in terms of a combination of two consecutive quaternary transitions named twisting and blooming ^2^. In this view, the intermediate conformation(s) illuminated by VCF corresponds to the completion of the un-blooming isomerization (i.e., compaction of the ECD) in a still closed-channel receptor. The apparent absence of significant twisting during GlyR activation (Figure S6) and its role on gating remain to be understood. Altogether, while the detailed activation mechanism may differ between different pLGIC subtypes, the collected data highlight a general activation mechanism in pLGICs characterized by a progressive conformational propagation from the ECD to the TMD.

### 4 Conclusion

This work illustrates the interest of combining VCF and MD approaches to identify functional intermediate conformations, characterize their flexibility, and annotate Cryo-EM structures to functional states. Noteworthy, such intermediate conformations may contribute to signal transduction in the postsynaptic membrane. The progressive transition also illuminates the mode of action of allosteric effectors and may be valuable for drug-design purposes.

## Materials and methods

### Site-directed mutagenesis

The gene coding the full-length human glycine ⍰1-subunit was cloned into the pMT3 plasmid containing an Ampicillin resistance gene. The PCR reaction was done with CloneAmp Hifi premix from Takara.

### ⍰1 homomeric glycine receptor expression in oocytes

*Xenopus Laevis* oocytes at stage VI are ordered from Portsmouth European Xenopus resource center and Ecocyte Biosciences and kept in Barth’s solution (87.34 mM NaCl, 1 mM KCl, 0.66 mM CaNO_3_, 0.72 mM CaCl_2_, 0.82 mM MgSO_4_ 2.4 mM NaHCO_3_, 10 mM HEPES and pH adjusted at 7.6 with NaOH). Ovary fragments obtained from Portsmouth European Xenopus resource center are treated as previously described ^51^. cDNA coding the ⍰1-subunit at 80 ng/μL is co-injected with a cDNA coding for GFP at 25 ng/μL into the oocyte nucleus by air injection. The injection pipettes are done with the capillary pipette (Nanoject II, Drummond) and PC-10 dual stage glass micropipette puller. Injected oocytes are incubated at 18°C for 3-4 days for expression.

### Labeling of the cysteine

MTS-TAMRA (Clinisciences) is dissolved in DMSO to obtain a final stock concentration of 10 mM and conserved at −20 °C. For labeling, oocytes expressing the receptors are incubated in 10 μM MTS-TAMRA (diluted in Barth’s solution to obtain a final DMSO concentration of 0.1%) at 16-18 °C for 20 mins before the recording. The procedure is similar for other fluorescent dyes (MTS-5-TAMRA, MTS-6-TAMRA).

### Two electrode voltage-clamp on oocytes

Oocytes expressing GlyR constructs are recorded in ND96 solution (96 mM NaCl, 2 mM KCl, 5 mM HEPES, 1 mM MgCl_2_, 1.8 mM CaCl_2_ and pH adjusted at 7.6 with NaOH). Molecules that are not 100% soluble in aqueous solution are first dissolved in DMSO and then diluted in ND96 to have a DMSO final concentration < 0.1 %. Currents were recorded by a Warner OC-725C amplifier and digitized by Digidata 1550A and Clampex 10 (Molecular Devices). Currents were sampled at 500 Hz and filtered at 100 Hz. The voltage clamp is maintained at −60 mV at room temperature. Recording pipettes were made with Borosilicate glass with filament (BF150-110-7.5, WPI) with a PC-10 dual stage glass micropipette puller to obtain pipettes of resistances comprised between 0.2 and 2 MΩ.

### Voltage-Clamp Fluorometry on oocytes

Injected oocytes are placed on a home-designed chamber with the animal pole turning toward the microscope objective. Currents were recorded by GeneClamp 500 voltage patch clamp amplifier (Axon instruments) and digitized by Axon Digidata 1400A digitizer and Clampex 10.6 software (Molecular Devices). TAMRA excitation is done by illumination with a 550 nm LED (pE-4000 CoolLED, 15%-20% intensity) through a FF01-543_22 bandpass filter (Semrock). The fluorescence emission goes through a FF01-593_40 bandpass filter (Semrock) and is collected by a photo-multiplicator H10722 series (HAMAMATSU). Recording pipettes were the same as those prepared for TEVC. Currents were sampled at 2 kHz and filtered at 500 Hz. The voltage clamp is maintained at −60 mV and at room temperature. The recording chamber has been designed to perfuse only the part of the oocyte from which the fluorescence emission is simultaneously collected (Figure S3). This chamber has several advantages over chambers that allow to record the current of the entire membrane of the oocytes: 1/ approximately the same population of receptors is recorded in current and fluorescence simultaneously, 2/ the chamber allows for an overexpression of the receptor on the oocyte membrane to increase the fluorescence signal without generating huge currents that make the clamp impossible to maintain during a prolonged period. The oocyte is placed on a hole and the geometry of the perfusion channels has been designed to “suck” the oocyte and stick it to the hole through a Venturi effect, ensuring efficient sealing between the two compartments. A limitation of the chamber is that the design is compatible with only relatively slow perfusion of the agonist, rapid perfusion often expelling the oocyte out of the hole.

### Voltage-Clamp Fluorometry data analysis

For each construct, we performed the experiments at least on 2 different batches of oocytes (oocytes from different ovary fragments and different animals) to obtained at least n=5. Data are analyzed by Clampfit software (Molecular Devices) and are filtered by a boxcar filter. Baseline was corrected from leaking currents and measurements were done to the peak of each response. Dose-response curves were fitted to the Hill equation by Prism Graphpad software, and the error bars represent the SEM values. Risetime and decaytime analysis were calculated in Clampfit by monoexponential fits of the onset and the offset of the recordings. Statistical analyses were done using a Student t-test in Prism Graphpad.

### Expression in cultured cells

Human Embryonic Kidney 293 (HEK-293) cells were cultured in Dulbecco’s modified Eagle’s medium (DMEM) with 10% FBS (Invitrogen) in an incubator at 37°C and 5% CO_2_. After being PBS washed, trypsin-treated (Trypsine-EDTA; ThermoFisher Scientific) and seeded on petri dishes, cells were transiently transfected using calcium phosphate-DNA co-precipitation with glycine receptor constructs (2 μg DNA) and a construct coding for a green fluorescent protein (0.2 μg). One day after transfection, cells were washed with fresh medium and recordings were carried out within 24 hours.

### Outside-out recordings and analysis

Recording currents are obtained with a RK-400 amplifier (BioLogic) using pClamp 10.5 software, digitized with a 1550 digidata (Axon instruments). Recording pipettes were obtained from thick-wall borosilicate glass (1.5X0.75mmx7.5cm, Sutter Instrument) using a micropipettes puller (P-1000, Sutter Instrument) and fire-polished with a micro-forge (MF-830, Narishige) to be used at resistances between 7 and 15 MΩ. Micro-pipettes were filled with internal solution (that contain in mM: 152 NaCl, 1 MgCl_2_, 10 BAPTA, 10 HEPES; pH adjusted to 7.3 with NaOH solution, osmolarity measured at 335 mOsm). Extracellular solution (in mM: 152 NaCl, 1 MgCl_2_, 10 HEPES; pH adjusted to 7.3 with NaOH solution and osmolarity was adjusted to 340 mOsm with glucose) was delivered by an automated perfusion system (RSC-200, BioLogic). Agonists’ solutions are freshly made before sessions of recordings and are obtained with extracellular solution added with 1 to 100 μM of glycine (dissolved from stock solution of 1M in water). Acquisition of recordings was performed at the sampling of 20 kHz and low-pass filtered at 1 kHz (using the amplifier 5-pole Bessel filter). Openings are analyzed using Clampfit 10 software and currents were calculated by fitting the all-points histogram distributions of current amplitudes with the sum of two gaussians curves. No further filtering is performed for the analysis.

### Molecular Dynamics simulations

For all the studied systems, i.e., apo-closed (PDB:6PXD), tau-closed (PDB:6PM3) and tau-open (PDB:6PM2), the protein was protonated at pH 7.0 using the Adaptive Poisson-Boltzmann Solver ^52^ via the webserver (https://server.poissonboltzmann.org/pdb2pqr). The protonated structures were then embedded in a POPC bilayer and solvated with TIP3P water molecules using CHARMM-GUI Membrane Builder ^53^. The CHARMM-36m forcefield ^54^ for the protein and CGenFF ^55^ for the taurine and strychnine were used for the all-atom molecular dynamics simulations. The simulations were performed using GROMACS 2021.4 ^56^ with input files generated by CHARMM-GUI Membrane Builder for the minimization and equilibration. In short, 5000 steps of steepest descent were performed, then the system was equilibrated over 6 short simulations in presence of atomic position restrains of decreasing strength, first in NVT, then in NPT ensemble, with the Berendsen thermostat and barostat. Finally, the equilibrated systems were carried on in production for 100ns at a 2fs timestep, in the NPT ensemble using V-rescale and Parrinello-Rahman as thermostat and barostat, respectively. In all cases, the LINCS algorithm was used to constrains the H-bonds and the PME served to treat the long-range Coulomb interactions. For each system we produced 5 replicas generated with different initial velocities from the equilibration.

### Trajectory analysis

The generated trajectories were analyzed using MDanalysis^57^ through Python scripts. Notably we computed the RMSD of the backbone atoms in the ECD, the TMD and the structural core (i.e., the inner and outer beta-sheets in ECD, and the M1-M3 helices in TMD) relative to the initial coordinates from cryo-EM. Additionally, we computed the RMSF decomposed per residue and averaged over the 5 subunits of the receptor. The plots were produced using Matplotlib ^58^, and the visual representations of the protein with the open-source version of PyMOL ^59^ (Schrödinger).

### Docking

To explore strychnine binding to the various conformational states of the receptor, i.e., apo-closed, tau-closed, and tau-open, we extracted 1 snapshot per nanosecond from the trajectory generated by Molecular Dynamics. By doing so, we ended up with 5 (replica) * 100 (ns) = 500 (snapshots) for each system. The molecular docking of strychnine was then executed using QuickVina 2.1 ^60^ with an exhaustiveness of 8 and a box of dimension 15*15*15Å centered on the previously defined binding sites. For this purpose, we prepared the PDBQT inputs of the receptor using *prepare_receptort4.py* script from MGLTools ^61^. Then, the positions of the 5 orthosteric binding sites of a given structure were computed on the fly as the center-of-mass of residues PHE223 of the (+)-subunit and PHE79 of the (-)-subunit using MDanalysis. Of note, for both tau-open and tau-closed, the receptor was simulated by Molecular Dynamics in the presence of taurine, but the docking of Strychnine was carried out without taurine.

## Acknowledgements

The work was supported the ERC (Grant no. 788974, Dynacotine), by the ‘Agence Nationale de la Recherche’ (Grant ANR-18-CE11-0015-01, Pentacontrol), Specific Grant Agreement No. 945539 (Human Brain Project SGA3), the doctoral school ED3C and the Foundation pour la Recherche Médicale (to Solène N. Lefebvre). The authors would like to thank Antoine Taly for help in data interpretation, and Alexandre Mourot for critical reading of the manuscript.

## Authors contribution

Sophie Shi and Solène N. Lefebvre set up and performed VCF experiments.

Laurie Peverini performed single-channel experiments and some TEVC experiments. Marc Gielen designed and constructed the recording chamber.

Adrien Cerdan performed MD and docking experiments.

Sophie Shi, Solène N. Lefebvre, Laurie Peverini, Adrien Cerdan, Marc Gielen, Marco Cecchini and Pierre-Jean Corringer designed the study and analyzed the data.

Sophie Shi, Laurie Peverini, Adrien Cerdan, Marco Cecchini, Pierre-Jean Corringer and Jean-Pierre Changeux wrote the manuscript.

Marco Cecchini, Pierre-Jean Corringer and Jean-Pierre acquired funding.

## Supplementary Figures

**Figure S1.**
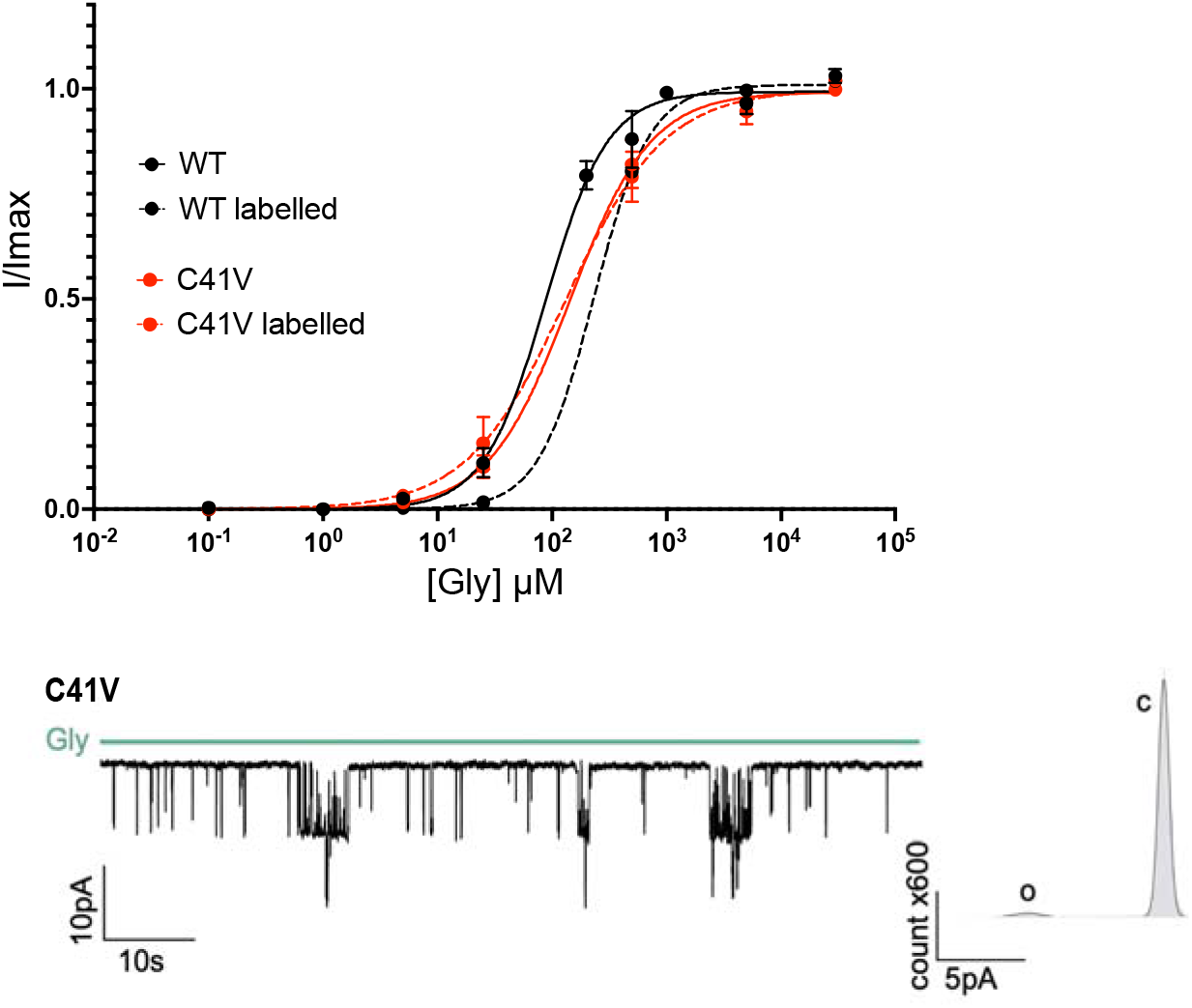
Control experiments showing labeled and unlabeled WT and C41V background mutant. Upper panel: current dose-response curves with mean ± the S.E.M. of the WT (black) and C41V (red) mutant without labeling in solid line and with MTS-TAMRA labeling in dotted line (n=4). The labeling of the WT triggers a weak loss of function of the receptor, while the C41V mutant is not impacted by the labeling. Lower left panel: single channel recording obtained by outside-out patch-clamp on HEK293 cells of the C41V mutant evoked by glycine application. The C41V mutant displays wild-type like glycine-elicited dose-response curve (TEVC) and unitary conductance as studied by outside-out single channel recording (Bormann et al., 1987 and 1993). Lower right panel: histograms of probability of opening showing the open (o) and the closed (c) state.

**Figure S2.**
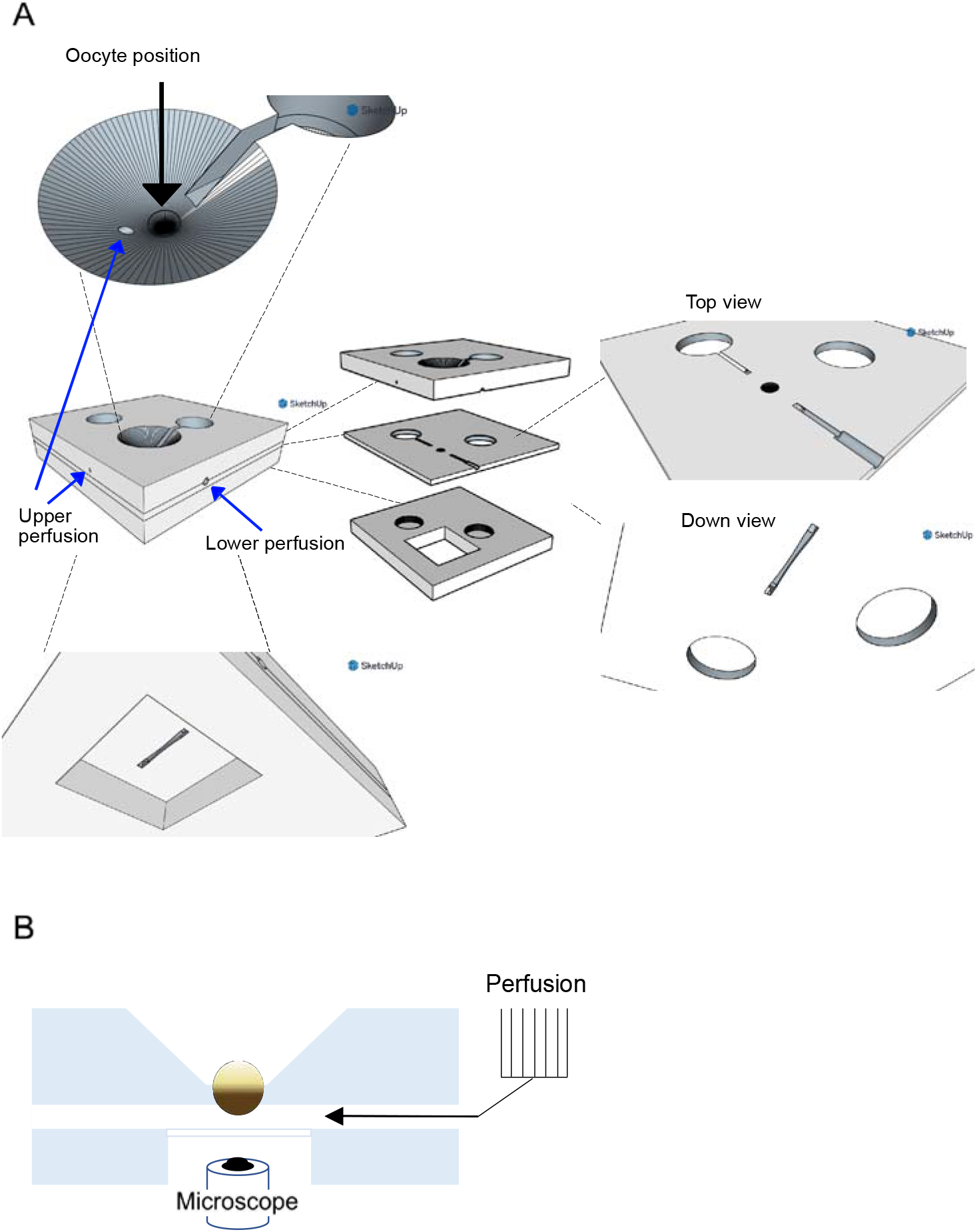
Blueprint of the perfusion recording chamber and schematic view of the voltage-clamp fluorometry set-up used in this study. (**A)** The recording chamber has been designed to perfuse only the portion of the animal pole that is imaged by fluorescence. A venturi effect allows to seal the oocyte in the chamber without any activation of receptors expressed on the upper side of the oocyte. **(B)** Schematic view of the voltage-clamp fluorometry set-up used in this study.

**Figure S3.**
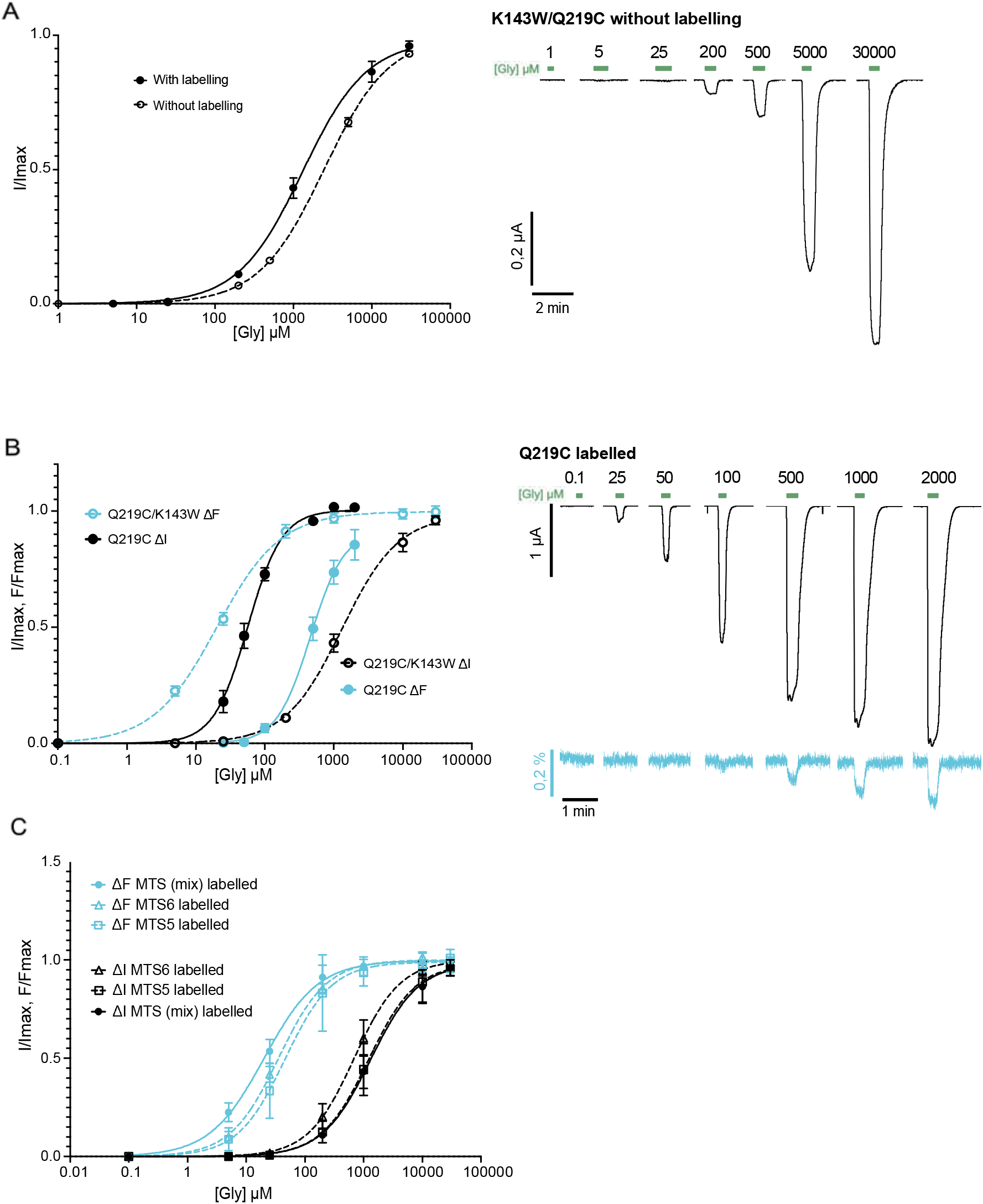
Control experiments showing labeled Q219C/C41V, unlabeled K143W/Q219C/ C41V, and MTS-5/6-TAMRA VCF data. **(A)** Left panel: dose-response curves with mean ± the S.E.M. of K143W/Q219C/C41V labeled with MTS-TAMRA (solid line) and unlabeled (dotted line, n=5). The labeling of the receptors triggers a small gain-of-function. Right panel: representative current recording of K143W/Q219C/C41V without labeling. **(B)** Left panel: VCF glycine dose-response curves with mean ± the S.E.M. of Q219C/C41V labeled with MTS-TAMRA (full line, n=6) compared to the K143W/Q219C/C41V (dotted line). For Q219C/C41V, the fluorescence (cyan) is shifted to a higher concentration of glycine compared to the current variation (black), with an EC_50_^fluo^ that is 10-fold higher than the EC_50_^current^. Right panel: representative VCF recordings. Fluorescence variation reaches 1,97 ± 0.49 % of ΔF/Fmax (n=6). **(C)** Fluorescence (cyan) and current (black) dose-response curves with mean ± the S.E.M. of mutant K143W/Q219C/C41V labeled with isomers of MTS-TAMRA (full line, round point), MTS-6-TAMRA (dotted line, triangular point, n=5) and MTS-5-TAMRA (dotted line, rectangular point, n=5).

**Figure S4.**
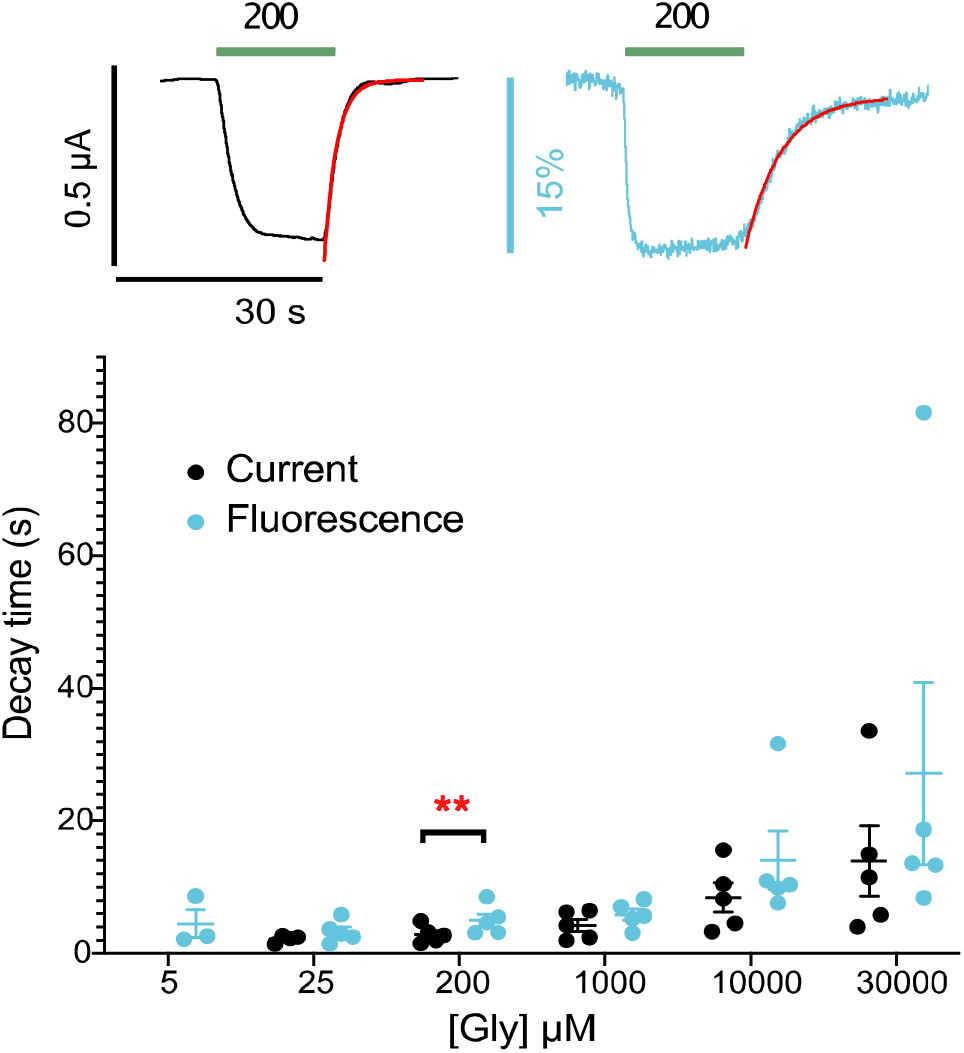
Single exponential fitting of the current and fluorescence traces offset of K143W/Q219C/C41V. Upper panel: single exponential fitting (red line) of the current and fluorescence traces offset. Lower panel: time constants τ (offset) values obtained via single exponential fitting with mean and S.E.M. error bars in second at different glycine concentrations. Unpaired student t-test revealed the significance of the difference between fluorescence and current offset (**: *P* < 0,005; ***: *P* < 0,0005).

**Figure S5.**
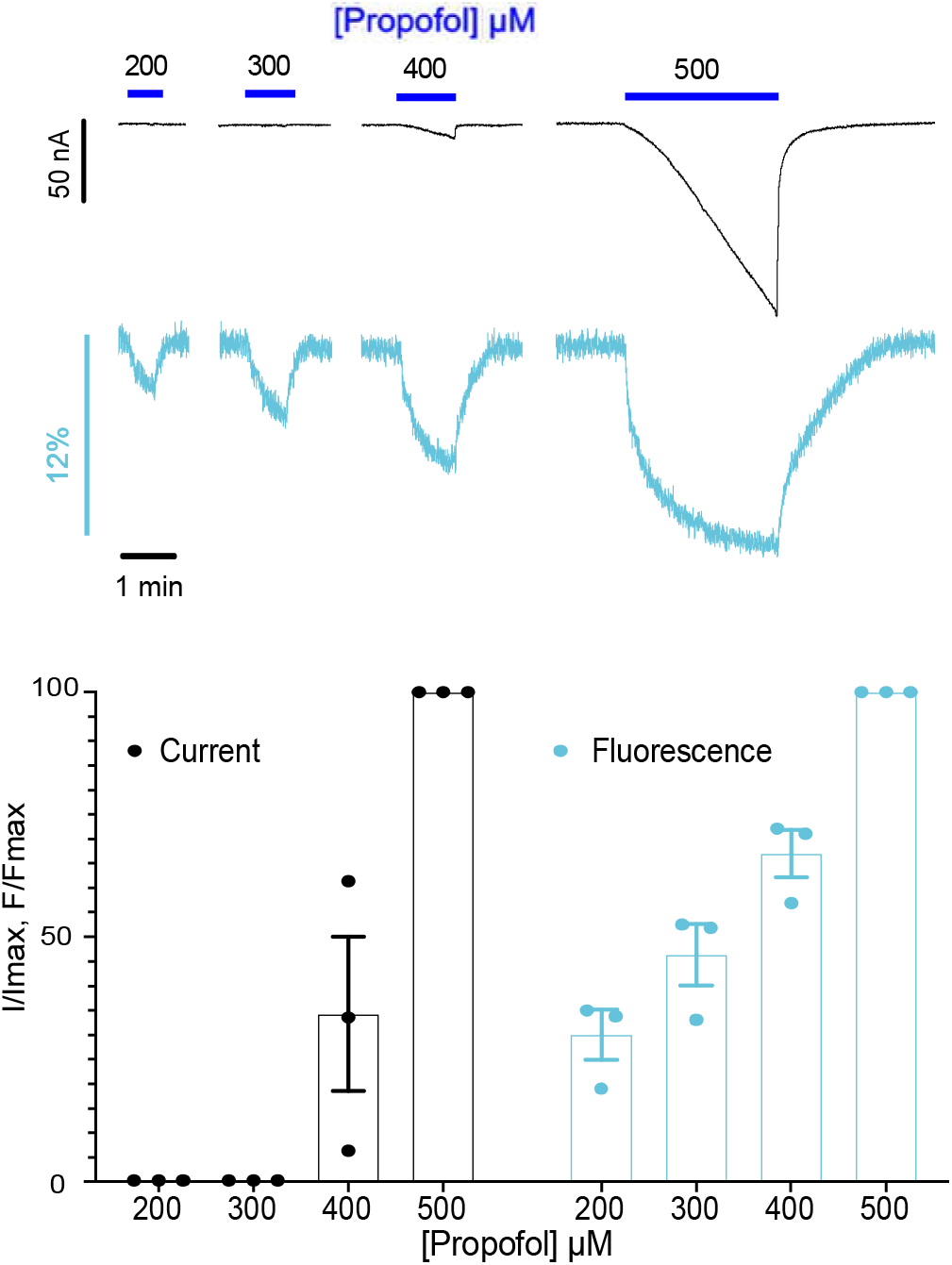
Electrophysiological and fluorescence characterization of propofol on K143W/Q219C/C41V. Upper panel: representative VCF recording under different concentration (200 to 500 μM) of propofol application. Lower panel: fluorescence (cyan) and current (black) variations normalized to the fluorescence and current variations under 500 μM of propofol (mean and S.E.M. (n=3). At low concentrations of propofol (less than 300 μM), a fluorescence variation is observed without any current. Of note, the slow kinetics of the fluorescence and current traces is due to the slow kinetics of partition of propofol into the plasma membrane (Heusser et al., 2018).

**Figure S6.**
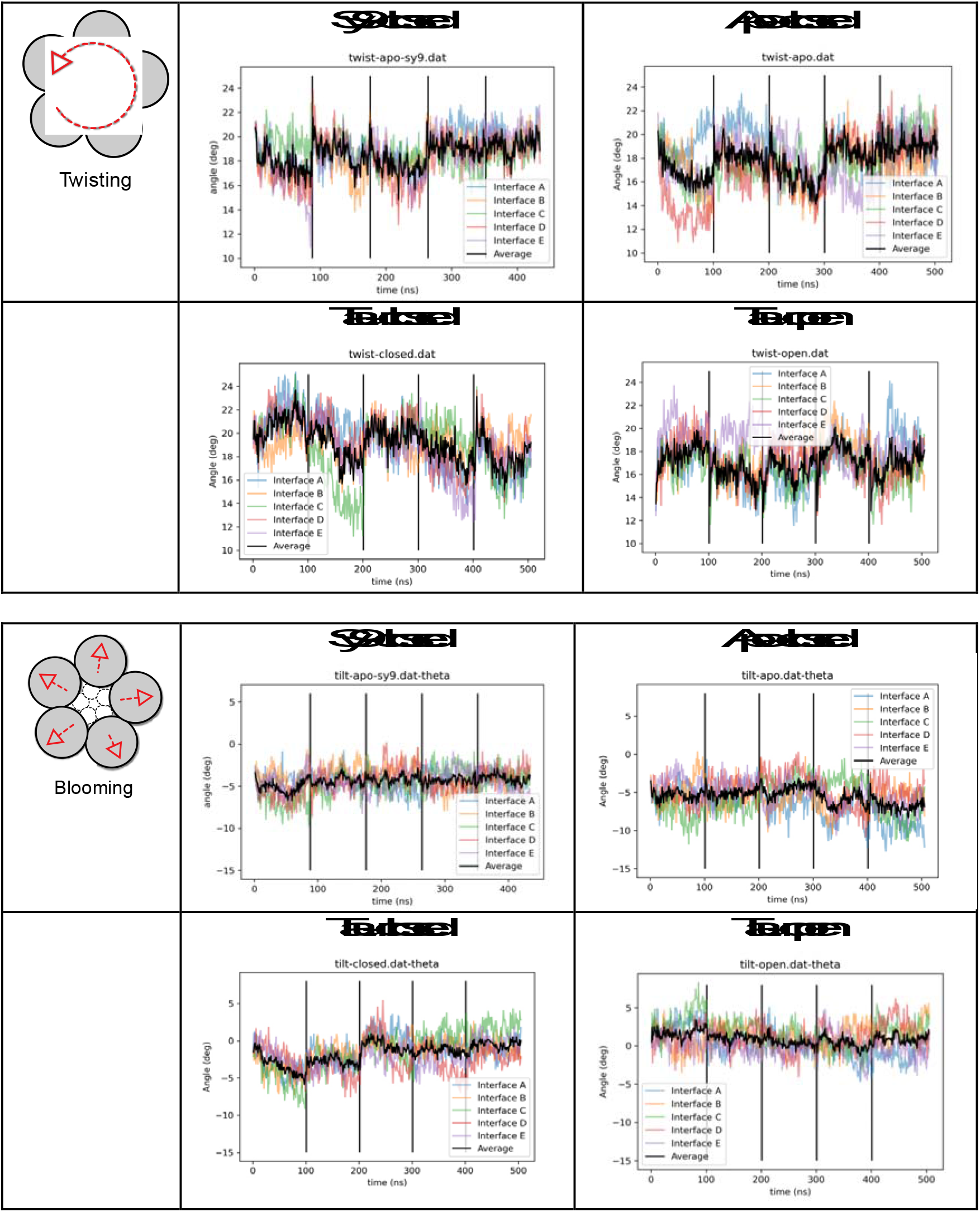
Twisting and blooming angles variations in the MD simulations. The four upper panels show the twisting angles, and the four lower panels the blooming angles for each set of simulation (apo-closed, stry-closed, tau-closed and tau-open). Each set of simulation corresponds to five independent runs that are shown separated by a vertical line. Twisting and blooming angles were computed as described in Martin et al., 2017. Color-coded lines correspond to values for individual subunits, the black thick line is an average over the pentamer. Concerning the twisting angle (top), no clear trend is observed. The average twisting angle fluctuates from 16 to 23 deg in the closed-channel conformations and is slightly lower in tau-open. The blooming angle (bottom) is more informative. Its average value is stable around 1 deg in tau-open, while it increases significantly in absolute value in stry-closed and Apo-closed around −5 deg. In tau-closed, blooming is more pronounced in replicas 1 and 2 where the RMSD from the initial cryo-EM coordinates is also the highest due to three unbinding events of taurine, while it remains close to 0 deg in the other replicas. Overall, this analysis suggests that receptor’s un-blooming, i.e., compaction of the ECD, is associated to agonist binding, as seen in apo-closed with tau-closed and tau-open. Additionally, an increase of the blooming angle in tau-closed is observed as a consequence of taurine unbinding. Finally, the active state represented by tau-open features a stable un-bloomed conformation.

**Table S1.** EC_50_ values for current and fluorescence responses to β-alanine and taurine at labeled K143W/Q219C/C41V mutant.

| Q219C/K143W C41V | EC<sub>50</sub><sup>current</sup> (μM) | n<sub>H</sub> | EC<sub>50</sub><sup>fluo</sup> (μM) | n<sub>H</sub> | n |
| --- | --- | --- | --- | --- | --- |
| Taurine | 7325.35 ± 2442.41 | 0.78 ± 0.11 | 104.46 ± 18.28 | 0.88 ± 0.05 | 6 |
| β-alanine | 1615.78 ± 502.42 | 1.22 ± 0.39 | 24.51 ± 5.96 | 0.83 ± 0.10 | 6 |

**Table S2.** ΔFmax/F (%) and ΔFmax/ΔImax (%) values to glycine for GlyR⍹1 mutants.

|  | ΔFmax/F (%) | ΔFmax/ΔImax (%) |
| --- | --- | --- |
| Q219C C41V | 1.97 ± 0.21 | 0.21 ± 0.03 |
| Q219C/K143W C41V | 12.2 ± 1.03 | 1.27 ± 0.43 |
| Q219C/K143W/N46D/N61D C41V | 15.19 ± 5.14 | 4.15 ± 0.76 |

**Table S3.**
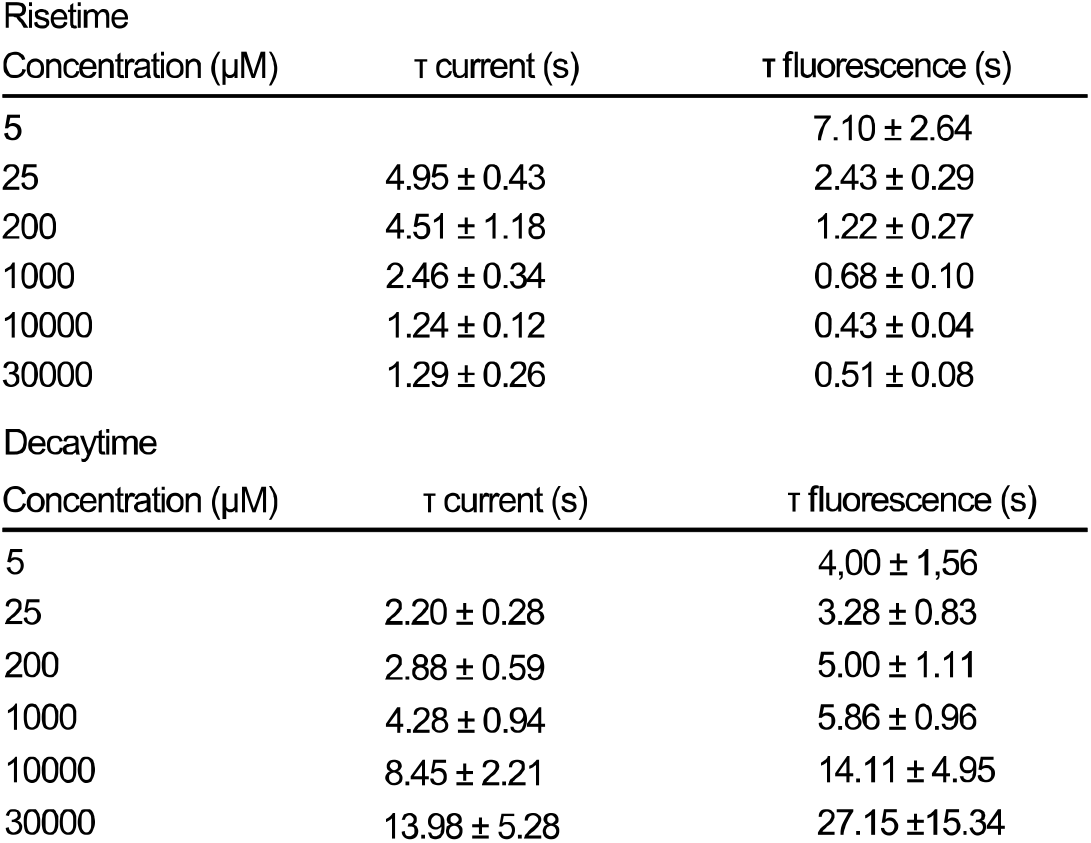
τ _rise_ and τ _decay_ values for fluorescence and current at labeled K143W/Q219C mutant.

